# Muc5ac mediates anti-viral immunity and virus-induced parasympathetic nerve dysfunction

**DOI:** 10.64898/2026.04.10.717757

**Authors:** James M. Kornfield, Shelby T. Hoffmeister, Ubaldo De La Torre, Chadwick B. Smith, Becky J. Proskocil-Chen, Christopher M. Evans, David B. Jacoby, Allison D. Fryer, Matthew G. Drake

**Author notes:** **Corresponding Author:** James Kornfield M.D. Division of Pulmonary, Allergy and Critical Care Medicine Oregon Health & Science University 3181 SW Sam Jackson Park Rd, Portland, Oregon 97239, United States.

## Abstract

Respiratory viruses can induce excessive bronchoconstriction in both asthmatic and healthy airways. Airway mucins such as Muc5ac form the first line of defense against inhaled pathogens. However, when produced in excess, they can also contribute to airway narrowing and mucus plug formation in asthma. In this study, we investigated the role of airway mucins in host defense against parainfluenza virus and in virus-induced airway hyperresponsiveness using Muc5ac-deficient (Muc5ac−/−) C57BL/6 mice. Parainfluenza virus infection induced airway hyperresponsiveness to inhaled methacholine in wild-type mice, an effect that was abolished in Muc5ac−/− mice. Parainfluenza virus–induced airway hyperresponsiveness was reversed by vagotomy, demonstrating it is mediated by parasympathetic nerve dysfunction. Muc5ac−/− mice exhibited higher viral titers, increased bronchoalveolar lavage cellularity, and elevated antiviral cytokine levels, but did not develop airway hyperresponsiveness. We did not see mucus plugging in any of our animals. Together, these findings indicate that Muc5ac is important for host defense against parainfluenza virus but paradoxically is also required for virus-induced airway hyperresponsiveness.

## Introduction

Respiratory viruses cause airway hyperresponsiveness, in which excessive narrowing of the airway occurs in response to stimuli, in both healthy airways and in asthma. Despite their significant clinical impact, few therapies directly target the effects of respiratory viruses. Parainfluenza virus, for example, accounts for 11% of adult asthma exacerbations(1), yet lacks any preventative or targeted therapeutic strategy to limit viral infection and airway hyperresponsiveness.

Respiratory viruses cause airway hyperresponsiveness in part by disrupting neural control of bronchomotor tone(2). Vagal parasympathetic nerves provide the dominant control of bronchoconstriction by releasing acetylcholine onto airway smooth muscle, where it activates smooth muscle M3 muscarinic receptors to provoke contraction and airway narrowing. Acetylcholine also binds presynaptic inhibitory M2 muscarinic receptors, which serve as a negative feedback mechanism to limit further acetylcholine release and bronchoconstriction. Virus infections impair M2 receptor signaling, resulting in loss of inhibitory feedback and excess acetylcholine release (2–4). inflammatory mediators, including TNFα(5) and IL-1β(6), mediate M2 dysfunction in non-atopic models, whereas eosinophil major basic protein mediates virus-induced M2 dysfunction after allergen sensitization(7).

Airway mucus hypersecretion has also been implicated as contributing to airflow obstruction. In healthy airways, mucus is composed primarily of the secreted glycoprotein Muc5b, while Muc5ac is upregulated in response to virus infection(8). Type 2 cytokines, such as IL-13, stimulate Muc5ac production and increase mucus plugs(9–11). Accordingly, mucus plugging has been associated with worse lung function in eosinophilic asthma(12) and is present extensively in the lungs after fatal asthma(13).

In this study, we tested the hypothesis that virus-induced mucus plugging is required for virus-induced airway hyperresponsiveness in non allergic airways by measuring airway physiology, mucus plugging, and immune responses in C57BL/6 wild-type and transgenic Muc5ac-deficient (Muc5ac -/-) mice infected with parainfluenza virus type 1. We demonstrated that Muc5ac is required for virus-induced airway hyperresponsiveness. However, airway hyperresponsiveness was mediated entirely by virus-induced airway nerve dysfunction, not mucus plugging. Loss of Muc5ac resulted in increased viral titers while paradoxically protecting against development of neuronal dysfunction, suggesting Muc5ac’s antiviral role is balanced against unexpected effects on airway nerves.

## Methods

### Mice

All studies were performed with 8-14 week-old mice on a C57BL/6 background. Muc5ac -/- mice were obtained from the University of Colorado and generated as previously described(14). C57BL/6 control mice were purchased directly from Jackson Laboratories (Jackson Research Laboratories, Bar Harbor, Maine). Protocols were performed in accordance with National Institute of Health guidelines and approved by the Oregon Health & Science University Institutional Animal Care and Use Committee.

### Virus propagation

Parainfluenza virus-type 1 (Sendai, American Type Culture Collection, Manassas, VA) was propagated in rhesus monkey kidney cell (RMK, Viromed, Minneapolis, MN) monolayers in RPMI medium for 1 week at 34^0^ C with 95% air/5% CO_2_. Virus stocks were purified by centrifugation at 10,000xg for 10 minutes, followed by sucrose gradient purification (15%/60% w/v) for 90 minutes at 65,000xg at 4^0^ C. Virus was collected from the sucrose interface, sedimented by centrifugation at 40,000xg for 45 minutes, and resuspended in RPMI medium.

### Virus Titer

Virus stocks were titered by hemadsorption assay using RMK cell monolayers. RMK cells were infected with serial 10-fold dilutions of parainfluenza-containing culture supernatants in triplicate. After incubating for 7 days, infectious titers were quantified per the Reed and Muench method(15). TCID_50_ (tissue culture infectious dose) of virus was defined as the amount of stock required to infect 50% of RMK monolayers.

### Virus Infection

Mice were sedated with ketamine (100mg/kg ip) and xylazine (10mg/kg ip) and inoculated intratracheally with 5 µL of RPMI containing 10^6^ TCID_50_ /ml of parainfluenza virus using a small animal laryngoscope. Control animals received 5 µL of RPMI media without virus. Airway responsiveness, inflammation, lung viral content, and mucus plugging were assessed 4 days after inoculation.

### Airway Physiology

Mice were anesthetized with ketamine (100 mg/kg ip) and xylazine (10 mg/kg ip), tracheostomized, paralyzed with succinylcholine (20 mg/kg im), and ventilated (200μl tidal volume, 120 breaths/min) with a positive end-expiratory pressure of 2 cmH_2_O. Body temperature was maintained between 36°C and 37°C. Heart rate was monitored by electrocardiogram. Inspiratory flow was measured via pneumotachograph (MLT1L, ADInstruments) and pressure was continuously recorded throughout the respiratory cycle with a differential pressure transducer (MLT0670, ADInstruments). Data were recorded using a PowerLab 4/SP analog-to-digital converter and analyzed with the LabChart Pro software (ADInstruments).

Plateau pressures (P_plateau_) were measured during 10 consecutive inspiratory pauses (225 ms) performed before and 1 minute after administration of aerosolized saline (vehicle) and methacholine (10μl, 10-1000 mM, Sigma Aldrich), delivered via in-line nebulizer. Airway resistance was calculated as P_peak_ − P_plateau_/inspiratory flow (cmH_2_O·mL^−1^·s^−1^) and averaged over 5 inspiratory pauses. Change in airway resistance was represented as the difference between airway resistance after each dose of methacholine and airway resistance after saline. In some experiments, nerve reflex bronchoconstriction was assessed by measuring the increase in airway resistance in response to methacholine before and after vagotomy.

### Vagal Nerve Stimulation

Vagus nerves were isolated bilaterally, the proximal ends were crushed, and the distal ends were attached to platinum electrodes. Nerves were submerged in mineral oil to prevent drying. Guanethidine (5mg/kg i.p.) was administered 20 minutes before electrical stimulation to block sympathetic nerve input. Nerves were electrically stimulated at 5-20hz and 16V with a 2ms pulse duration. Pulse trains were run for 10 seconds at 70-second intervals. After a frequency response curve was measured, atropine (3mg/kg i.p.) was administered followed by electrical stimulation at 20hz 5 minutes later. Bronchoconstriction was expressed as the difference in peak pressure (Δpeak pressure) with electrical stimulation compared to the unstimulated baseline.

### Bronchoalveolar Leukocyte Quantification

Bronchoalveolar lavage (BAL) fluid was collected from the right lung by clamping the left mainstem bronchus to preserve left lung intraluminal mucus. Total leukocytes and differential cell counts were performed on BAL fluid using a hemocytometer (Hausser Scientific) and by manual counting following hematoxylin and eosin staining, respectively. Following lavage, right lungs were flash frozen in liquid nitrogen.

### Histologic Quantification of Mucus Plugging

Left lungs were fixed in methacarn for 16 hours, paraffin-embedded, cut into 10μm sections, and stained with Alcian Blue and Periodic Acid Schiff (AB-PAS, Abcam). Slides were scanned using a Nikon Eclipse CI upright microscope, and were examined by 2 separate blinded reviewers at 20X magnification using ImageJ. Nonaxial airways (defined by the absence of cartilage in the airway wall) were assessed in 5 non-sequential lung sections per animal.

### MicroCT Quantification of Mucus Plugging

MicroCT images were acquired using a Siemens Inveon microCT (80kV, 500 microamps, 0.5mm aluminum filter) post-mortem after inflating lungs to an airway pressure of 10cmH_2_0. Images comprised a 360-degree imaging pattern with 1 exposure per position and 0.5 degrees of arc separation between positions for a total of 720 positions with 1 sec settle time at each position. The calculated voxel size was 36 microns. Mucus plugging was measured using a modified Dunican score(12) with a composite of mucus plugs present (1) or absent (0) within each of the 4 lobes of the right lung and superior and inferior segments of the left lung.

### Real-time RT-PCR

RNA was isolated from homogenized lung using an RNeasy Mini Kit (Qiagen, Germantown, MD), cDNA was created using Superscript III Reverse Transcriptase (Invitrogen, Carlsbad, CA) and PCR was performed on a 7500 Fast RT PCR system (Applied Biosystems, Carlsbad, Ca). Parainfluenza RNA CT values were converted to TCID_50_/ml using a parainfluenza standard curve quantified by RMK cell titer as described above. Parainfluenza primers were custom ordered as follows: parainfluenza matrix protein 5’-ATGCGGCTGATCTTCTCACT-3’ and 5’-CTTTGCCACGACATTAGGGT-3’, parainfluenza fusion protein 5’-GAGTGGCAACATCAGCACAG-3’ and 5’-CTTATCGCGGGTTTGATCTC-3’, parainfluenza nucleocapsid protein 5’-GAGCTATGCAATGGGAGTCG-3’ and 5’-TTGCCAAATGATGTCTGAGC-3’. Interleukin 6, TNFα, IFNγ, and RT^2^ Profiler PCR Arrays were quantified using RT² qPCR Primer Assay for Mouse (Qiagen, Germantown, MD) according to the manufacturer’s instructions. Ct values were normalized to 18s ribosomal RNA levels and expressed as a fold change (2^-ΔΔCt^).

### Statistical Analyses

Two-way repeated measures ANOVA test was used to compare methacholine dose response curves and vagal nerve frequency response curves. One-way ANOVA with Tukey’s multiple comparisons test was used to compare group BAL cell count and differentials as well as Interleukin 6, TNFα, IFNγ RNA expression levels. Unpaired T-tests were used to compare viral RNA titers as well as histologic airway measurements. Data are graphed and analyzed using GraphPad Prism 10 and presented as mean ± SEM. P values < 0.05 were considered significant.

## Results

### Muc5ac is required for virus-induced airway hyperresponsiveness

Virus-infected WT mice had increased responsiveness to methacholine compared to mock-infected WT mice (**Figure 1A**). In contrast, virus-infected Muc5ac -/- mice had similar airway responsiveness to mock-infected Muc5ac-/- and mock-infected WT control mice (**Figure 1B**), indicating that Muc5ac is required for development of hyperresponsiveness. Deletion of Muc5ac did not change the response to methacholine. Lung compliance (**Figure 1C-D**) and baseline lung resistance before methacholine administration were similar across genotypes and infection status for all groups (**Figure 1E**).

**Figure 1.**
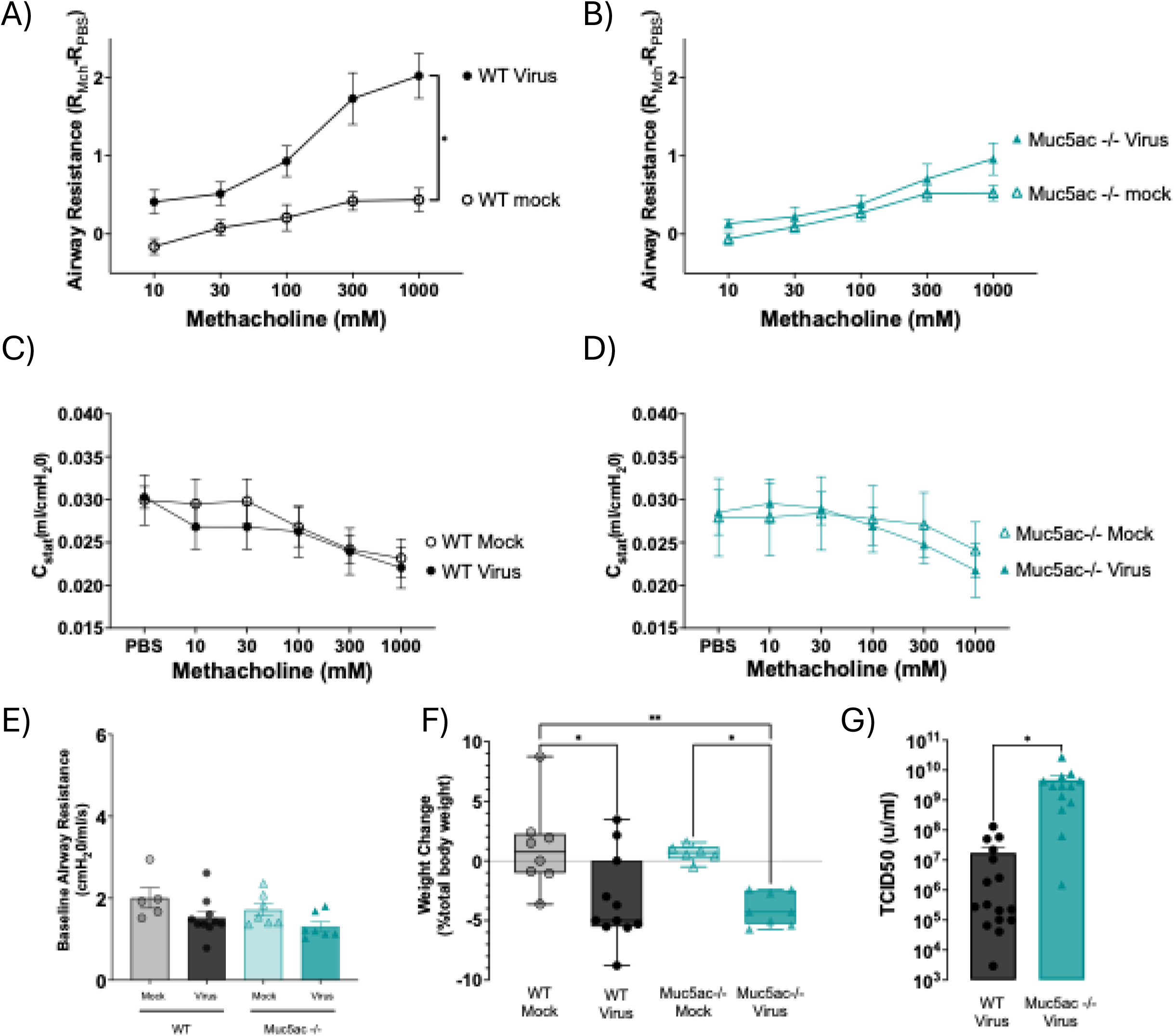
Muc5ac is required for parainfluenza virus-induced airway hyperresponsiveness and attenuates viral replication. Virus infection increased airway resistance in wild-type mice **(A)** but not Muc5ac -/- mice **(B).** Static lung compliance was similar in all groups (**C-D)**. Baseline airway resistance was similar between all groups **(E)**. Unlike mock-infected mice, virus-infected WT and Muc5ac -/-mice lost weight four days after inoculation **(F)**. Viral titers in lungs were increased in Muc5ac -/- mice compared to wild-type mice 4 days after inoculation **(G)**. n=5-16 mice per group. Group means +/- SEM are shown. \**p* < 0.05.

### Muc5ac decreases parainfluenza virus lung content

Parainfluenza virus infection caused weight loss in WT and in Muc5ac -/- mice, in contrast to weight gain in both mock infected WT mice and in mock-infected Muc5ac -/-mice (**Figure 1F**). Viral titers were significantly increased in Muc5ac -/- mice compared to WT mice (**Figure 1G**).

### Virus-induced airway hyperresponsiveness is mediated by parasympathetic nerve dysfunction

Airway responsiveness to inhaled methacholine was tested before and after bilateral surgical vagotomy to determine the role of nerve-mediated reflex bronchoconstriction in virus-induced airway hyperresponsiveness. Vagotomy eliminated airway hyperresponsiveness in virus-infected WT mice (**Figure 2A**). As before, Muc5ac -/- mice were protected from virus-induced airway hyperresponsiveness before vagotomy.

**Figure 2.**
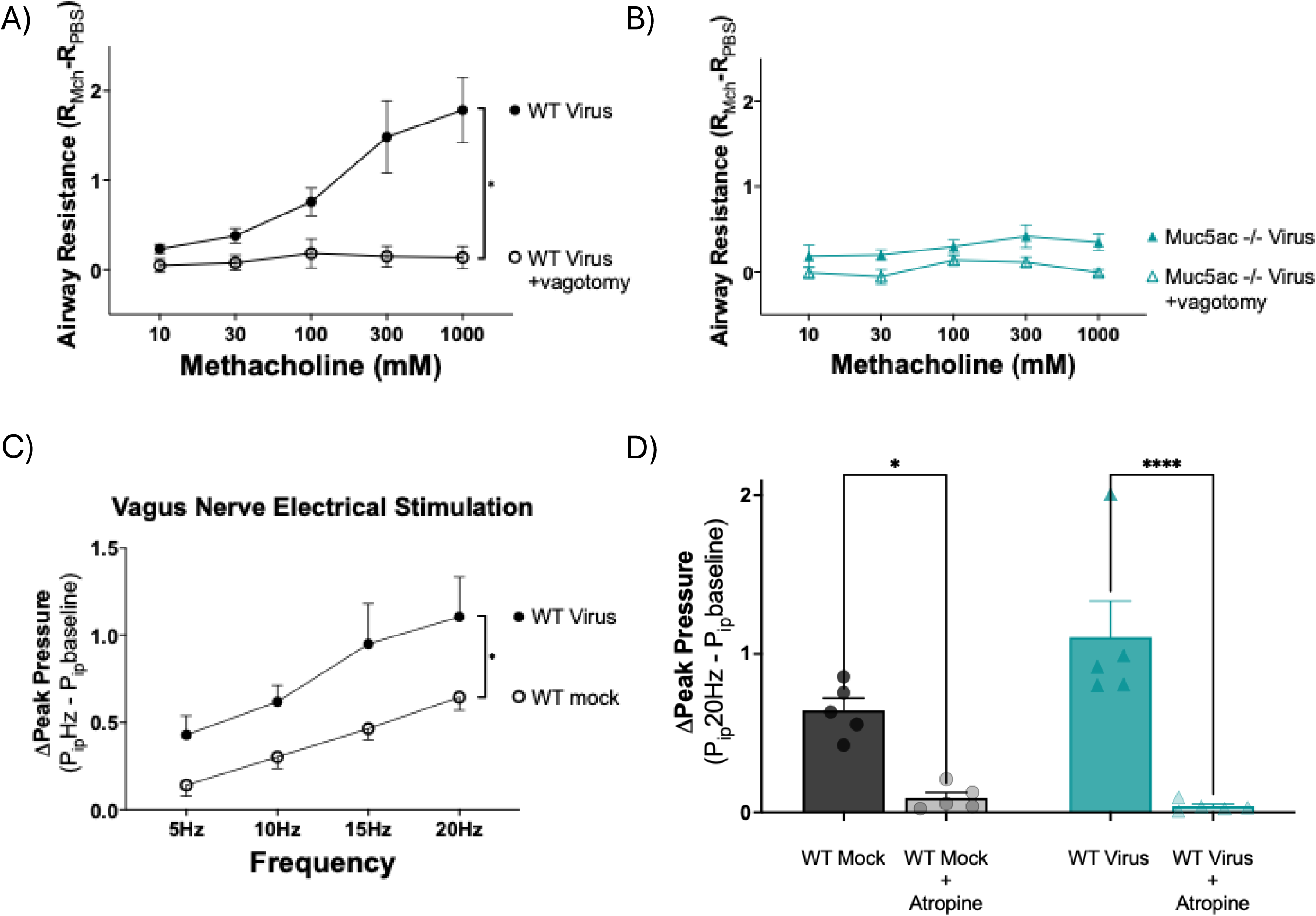
Parainfluenza virus-induced airway hyperresponsiveness is mediated by parasympathetic nerve dysfunction. Increased airway resistance in virus infected wild-type mice was eliminated by vagotomy **(A).** Virus-infected Muc5ac -/- mice did not developed increased airway resistance and vagotomy had not additional effect in this group (**B).** Bronchconstriction induced by electrical stimulation of the vagus nerves was increased in virus-infected wild-type mice **(C)**. Vagal nerve stimulation induced bronchoconstriction was blocked after the administration of atropine **(D)**. n=4 mice per group. Group means +/- SEM are shown. \**p* < 0.05 using a two-way ANOVA.

Vagotomy had no additional effect on virus-infected Muc5ac -/- mice (**Figure 2B**).

Next, we isolated and tested the role of efferent parasympathetic nerves in virus-induced airway hyperresponsiveness. Vagus nerves containing parasympathetic nerve fibers were crushed proximally, and the distal ends were electrically stimulated to provoke bronchoconstriction. In virus-infected WT mice, bronchoconstriction induced by electrical stimulation of the vagus nerves was significantly increased compared to mock-infected WT mice (**Figure 2C**). Bronchoconstriction in response to electrical stimulation was blocked after the administration of atropine (**Figure 2D**). Muc5ac -/- mice were not tested since they do not develop airway hyperresponsiveness after virus infection.

### Intraluminal mucus does not explain virus induced airway hyperresponsiveness

Airway diameter and intraluminal mucus content were quantified in methacarn-fixed lung sections stained with Alcian Blue – Periodic Acid Schiff (**Figure 3A-B**). Overall, mucus occlusion of airway lumens was similar between virus-infected WT and virus-infected Muc5ac -/- mice (**Figure 3C**) and likewise the contribution of mucus to airway wall thickness was similar between virus-infected WT and Muc5ac -/- mice (**Figure 3D**). Airway epithelial thickness was reduced in Muc5ac -/- compared to WT mice (**Figure 3E**), as was airway smooth muscle thickness (**Figure 3F**). Post-mortem microCT images similarly demonstrated minimal mucus plugging in virus-infected WT mice (**Figure 3G-H**).

**Figure 3.**
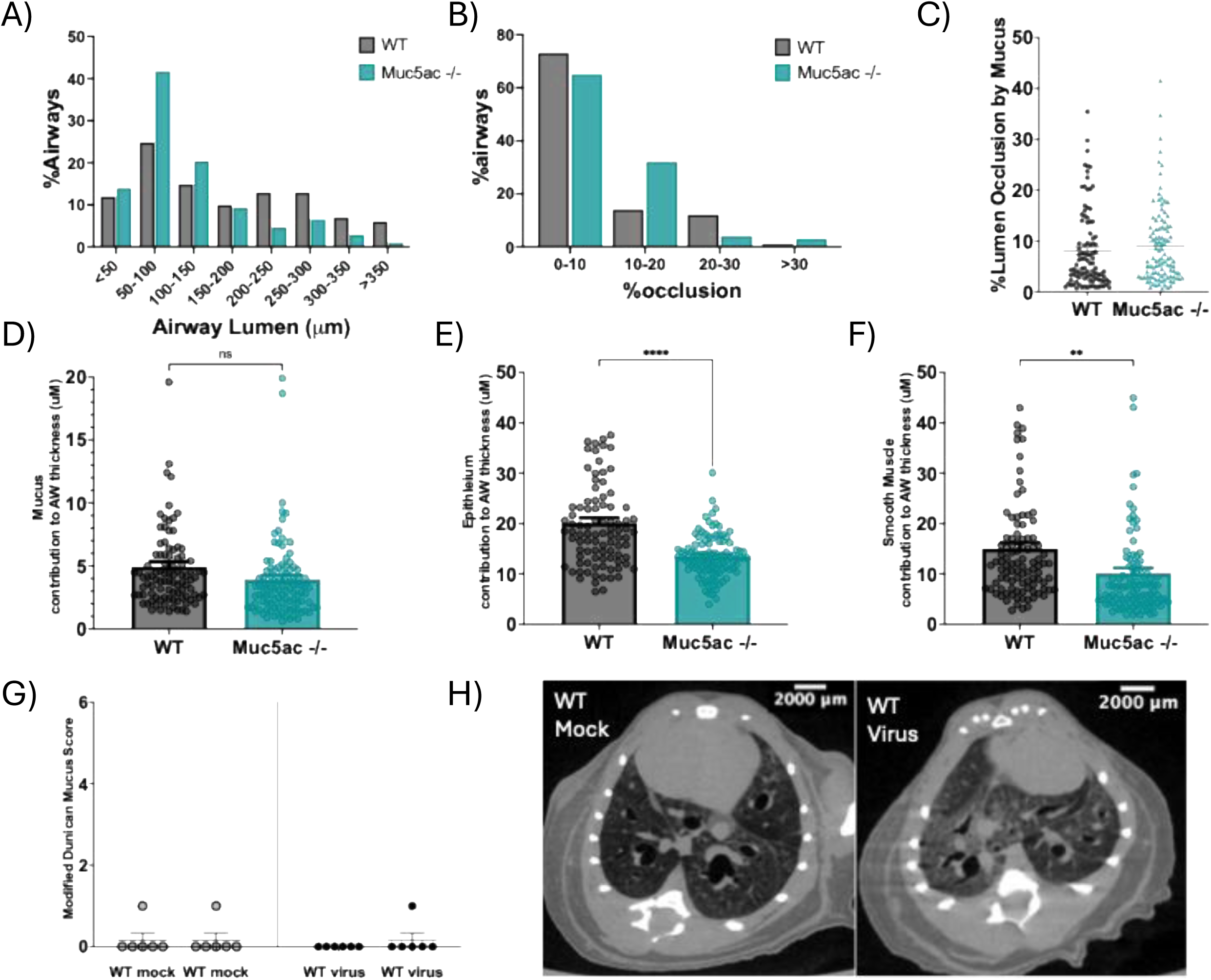
Intraluminal mucus accumulation was similar between virus-infected wild-type and Muc5ac -/- mice. Airway lumen diameter **(A)**, the proportion of airways with mucus occlusion **(B)** the percentage of mucus occlusion within each airway **(C)**, and airway wall thickness **(D)** were similar between parainfluenza virus-infected wild-type and Muc5ac -/- mice. Epithelial and smooth muscle thickness were decreased in Muc5ac -/- mice compared to WT mice **(E-F).** Mucus plugging was similar on MicroCT images in virus infected wild-type and Muc5ac -/- mice **(G-H).** Group means +/- SEM are shown. *p<0.05 using a T-test.

### Muc5ac -/-mice have increased airway cellular inflammation after virus infection

Total leukocytes were increased in bronchoalveolar lavage fluid from virus-infected Muc5ac-/- mice compared to virus-infected WT mice (**Figure 4A**). This increase was driven by a marked rise in bronchoalveolar lavage fluid neutrophils (**Figure 4B**).

**Figure 4.**
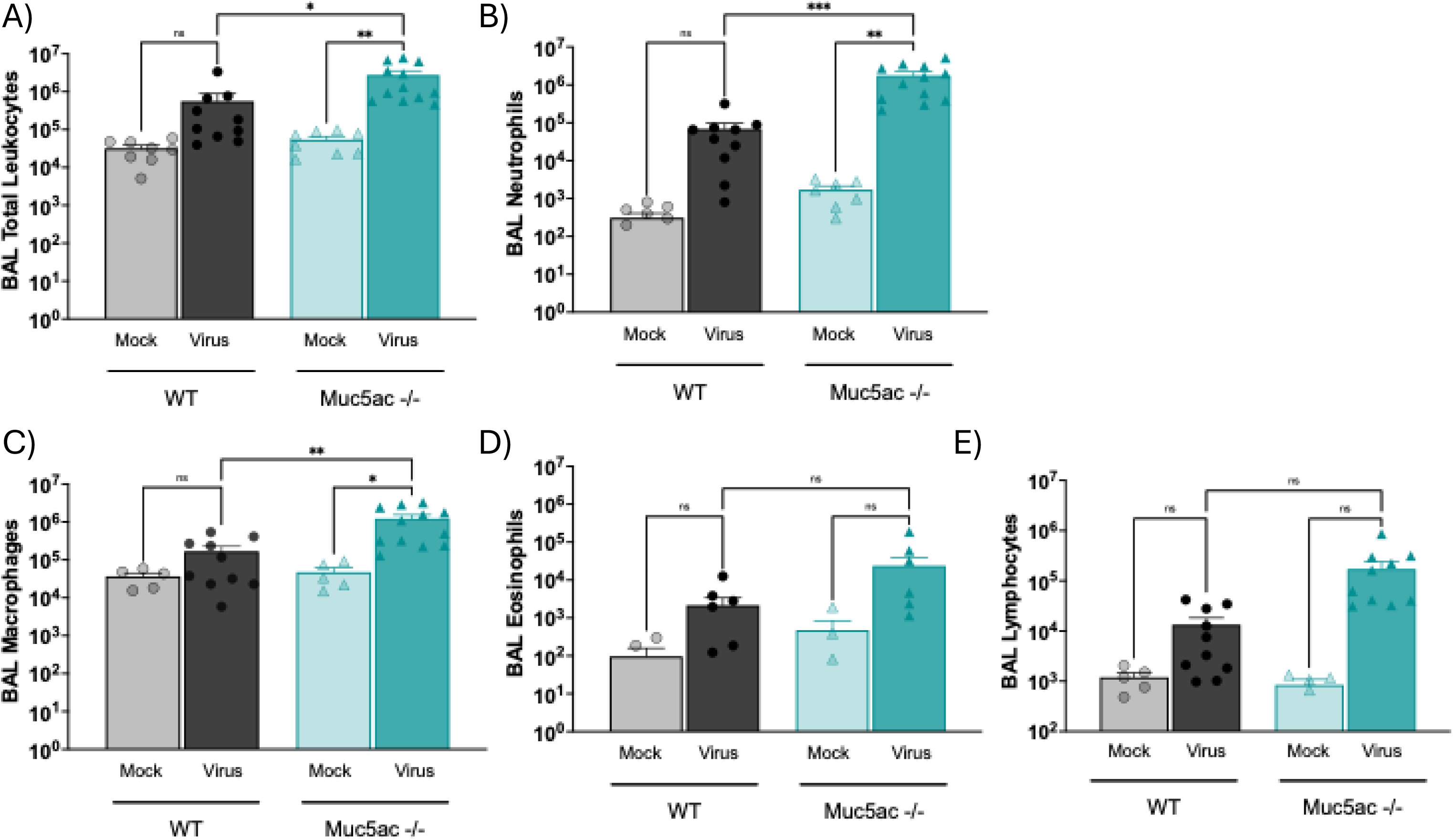
In the absence of Muc5ac leukocytes in bronchoalveolar lavage fluid were increased after virus infection. The total number of leukocytes in the bronchoalveolar lavage fluid was increased after parainfluenzaz virus infection in both WT (grey) and Muc5ac -/- mice (green) compared to mock controls and this effect was enhanced in Muc5ac-/- infected mice **(A).** The increased cellularity in lavage fluid of Muc5ac-/- mice was predominantly an increase in neutrophils **(B)** though macrophages were also significantly increased relative to WT infected mice **(C).** There was a trend toward increased eosinophils and lymphocytes in lavage fluid after infection though this did not reach statistical significance **(D-E).** n=6-12 mice per group. Group means +/- SEM are shown. \**p* < 0.05 using a one-way ANOVA.

Macrophages, lymphocytes, and eosinophils were also increased in Muc5ac-/- animals after parainfluenza virus infection compared to WT infected mice (**Figure 4C-E**).

### Anti-viral cytokines are increased after parainfluenza virus infection

Virus infection increased expression of IL-6, TNFα, and IFN-γ in WT mice compared to mock-infected controls. Cytokine transcription was further increased in virus-infected Muc5ac -/- mice compared to infected WT with a mean TNFα fold increase of 12.13, while there were increased in IFN-γ and IL-6 expression in Muc5ac -/- virus infected mice with mean fold change of 40.61, and 90.06 respectively (**Figure 5A-C**).

**Figure 5.**
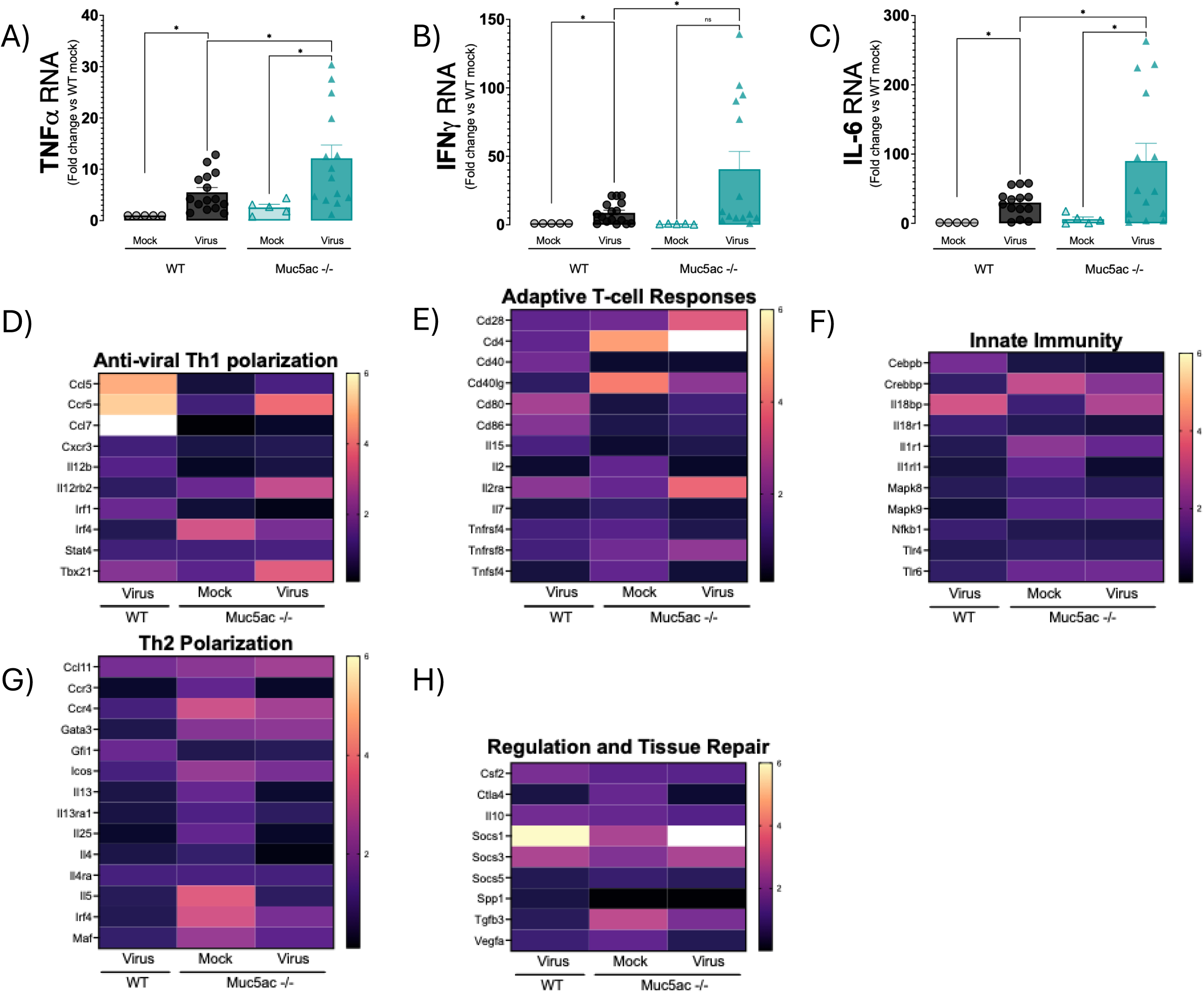
Muc5ac knock out lead to an amplified anti-viral cytokine response to Parainlfuenza but differential expression of chemokines. Virus infection caused a significant increase in TNFα, IFN-γ and IL-6 RNA expression measured by fold change using RT-PCR with Muc5ac -/- (green) demonstrating a potentiated response in transcription of these cytokines compared to WT mice (grey) **(A-C)** n=5-14 mice per group, group means +/- SEM are shown *p<0.05 using a one-way ANOVA. Qiagen RT^2^ profiler array was used to measure expression of host immune response genes which were categorized based on their physiologic roles. Gene expression was measured using fold change compared to WT mock infected controls and heat maps were generated on a scale of 0.1 – 6 **(D-G)** n=4 mice per group.

### Virus infection differentially regulates pulmonary chemokine expression in wild-type and Muc5ac mice

Cytokine and chemokine expression was quantified using an RT-PCR cytokine array. Virus infection significantly upregulated chemokines CCL5 (RANTES) and CCL7 (MCP-3), and CCR5 receptor, in WT mice compared to Muc5ac -/- mice and both mock-infected groups. In virus-infected Muc5ac -/- mice, CCL5 and CCL7 were similar to mock-infected controls. (**Figure 5D**).

Muc5ac -/- mice had increased expression of genes involved in the IFN-γ signaling cascade, including IL12rb2 and Tbx21 encoding the T-bet protein, which correlated with increased IFN-γ expression in Muc5ac -/- infected mice compared to WT mice (**Figure 5D**). T-cell signaling, specifically IL-2ra and CD4, was also differentially regulated in virus-infected Muc5ac -/- mice compared to WT mice. Uninfected Muc5ac -/- mice also had increased expression of CD4 and other T-cell activating genes at baseline (**Figure 5E**).

Innate immune activation markers were not elevated by day 4 after virus infection. Specifically, TLR 4, TLR 6, and NFκB1 were unchanged in infected mice, as were IL-1-associated genes, while counter-regulatory IL-18 binding protein expression was increased after virus infection in both Muc5ac -/- and WT mice (**Figure 5F**). Th2 cytokines, including IL-4, IL-5, and IL-13, and chemokines, such as CXCR3 and eotaxin-1 (CCL11), were unchanged four days after virus infection in both Muc5ac -/- and WT mice (**Figure 5G**). Finally, the counter-regulatory proteins SOCS1 and SOCS3 were increased 4 days after infection in both WT and Muc5ac -/- mice (**Figure 5H**).

## Discussion

Our results demonstrate that the airway mucin Muc5ac is required for the development of virus-induced airway hyperresponsiveness, mediated through airway parasympathetic nerves. Virus-incuded hyperresponsiveness is independent of airway mucus accumulation. We also show that Muc5ac serves a protective role in the defense against parainfluenza virus by limiting viral replication and airway inflammation. These findings uncouple viral load, host inflammatory responses, and mucus plugging from airway hyperresponsiveness.

Vagotomy completely abolished airway hyperresponsiveness in virus-infected WT mice, indicating that hyperresponsiveness is entirely vagally mediated. Since vagal electrical stimulation also produced exaggerated bronchoconstriction in virus-infected WT mice, we conclude that parasympathetic nerve dysfunction is a primary cause for excessive bronchoconstriction. Muc5ac-deficient mice were protected against virus-induced airway hyperresponsiveness and vagotomy had no additional effect, suggesting that Muc5ac is required for development of virus-induced neuronal dysfunction.

Respiratory viruses are known to cause airway hyperresponsiveness that is mediated by parasympathetic nerve dysfunction in guinea pigs(4–6,16–18) and rats(19). To our knowledge, our results are the first to show parasympathetic nerve dysfunction in mice using electrical vagal stimulation. Parasympathetic nerves release acetylcholine to activate M3 muscarinic receptors on airway smooth muscle to provoke bronchoconstriction. These nerves also express inhibitory presynaptic M2 muscarinic receptors, which reduce neuronal acetylcholine release to prevent excessive bronchoconstriction. In guinea pigs, TNF-α(5) and IL-1β(6) mediate M2 receptor dysfunction, resulting in loss of inhibitory feedback, increased acetylcholine release, and excessive bronchoconstriction. In our study, TNF-α, IL-6, and IFN-γ were markedly elevated in Muc5ac -/- mice, yet these mice did not develop airway hyperresponsiveness, suggesting cytokine production alone does not fully explain the mechanism of virus-induced parasympathetic nerve dysfunction.

Prior studies have established that Muc5ac plays an important role in the formation of the airway mucus plugging in both human asthma(12,20–22) and in animal models using allergen sensitization and challenge(23). In these studies, Muc5ac was required for mucus accumulation after allergen exposure, which, in conjunction with increased airway smooth muscle contraction, narrows airways and increases airway resistance(23). Our data suggest alternative mechanisms for airway resistance exist in the setting of parainfluenza virus-induced airway hyperresponsiveness, independent of mucus plugging or smooth muscle contractility. Overall, mucus plugging was similar between virus-infected WT and Muc5ac -/- mice, despite only WT mice developing airway hyperresponsiveness, and hyperresponsiveness was abolished by vagotomy, indicating that smooth muscle contractility was not a dominant mechanism in the absence of neuronal signaling. These observations support the conclusion that increased airway resistance after parainfluenza virus infection is mediated entirely by airway nerve dysfunction.

Our data suggest that Muc5ac plays a key role in host defense against parainfluenza virus infection, since the absence of Muc5ac resulted in significantly increased viral titers and exaggerated inflammatory cell recruitment. All leukocyte lineages were increased after virus infection, including, most prominently, neutrophils. Muc5ac may attenuate viral replication through multiple mechanisms. Intriguingly recent data suggest that virus may be trapped in the mucus layer by IgG allowing for mucociliary clearance of pathogens (24,25). Alternatively, Muc5ac sialic acid residues may also serve as decoy binding sites for viral hemagglutinin, a critical step in the attachment and uptake of viruses at airway epithelium(26,27). Despite higher viral titers and markedly increased inflammatory cell responses, however, Muc5ac -/- mice were protected from virus-induced airway hyperresponsiveness, suggesting that disrupting Muc5ac may have therapeutic potential in virus-induced bronchospasm.

Prior studies have demonstrated that the localization of immune cells to airway nerves is central in the development of neuronal dysfunction most notably eosinophils(28–31), mast cells(32–35) , and dendritic cells(11,36–38). Accordingly, we observed an increase in CCR5 expression and its ligand CCL5 in virus-infected WT mice, and saw a similar effect on CCL7. These chemokine signaling pathways drive inflammatory cell migration into the airways in RSV, rhinovirus, and influenza infections, and in the case of CCL5, have been associated with the development of airway hyperresponsiveness(39). We hypothesize that upregulation of the CCL5-CCR5 axis as well as CCL7, may enhance leukocyte trafficking to airway nerves and that this proximity may be necessary to promote nerve-mediated hyperresponsiveness. Moreover, since these chemokines were not upregulated in virus-infected Muc5ac -/- mice, Muc5ac may have an underappreciated immunomodulatory role in orchestrating the host immune response.

Recent evidence suggests that the gel-forming mucins, like Muc5ac, closely associate with soluble mediators in the airway through electrostatic and glycan interactions.

Myriad proteins bind to Muc5ac and can be sequestered in the mucin layer(40). For example, CXCR2 ligands have been shown to associate with Muc5ac in a glycosylation-dependent manner(41), supporting the idea that Muc5ac can spatially organize inflammatory mediators. Such interactions may allow Muc5ac to concentrate chemokines, shaping the spatial distribution of inflammatory cells and enhancing localized neuroimmune interactions that produce airway hyperresponsiveness.

Collectively, our findings suggest new roles for Muc5ac during viral respiratory infection. While Muc5ac limits viral replication and attenuates inflammation, it is also required for the development of virus-induced airway hyperresponsiveness through a parasympathetic nerve–dependent mechanism that occurs independently of mucus plugging, cytokine levels, or virus burden. These results have important implications for therapeutic strategies targeting mucus in virus-induced bronchoconstriction and asthma exacerbations. Interventions aimed at broadly suppressing mucin production may inadvertently impair host defense, while modulating neuro-immune interactions could offer a more targetted approach to preventing virus-induced bronchoconstriction.

